# Robust Stability of Multicomponent Membranes: the Role of Glycolipids

**DOI:** 10.1101/2019.12.19.882092

**Authors:** Yuan Chen, Arjen Doelman, Keith Promislow, Frits Veerman

## Abstract

We present the multicomponent functionalized free energies that characterize the low-energy packings of amphiphilic molecules within a membrane through a correspondence to connecting orbits within a reduced dynamical system. To each connecting orbits we associate a manifold of low energy membrane-type configurations parameterized by a large class of admissible interfaces. The normal coercivity of the manifolds is established through criteria depending solely on the structure of the associated connecting orbit. We present a class of examples that arise naturally from geometric singular perturbation techniques, focusing on a model that characterizes the stabilizing role of cholesterol-like glycolipids within phospholipid membranes.

## 1 Introduction

Amphiphilic molecules play a fundamental role in the self-assembly of nanostructured membranes. These include phospholipids, the building blocks of cellular membranes, and synthetic polymers that are finding applications to drug delivery compounds and as active materials for separator membranes in energy conversion devises, [5, 13, 25]. The scalar functionalized Cahn-Hilliard free energy models the interaction of a single species of amphiphilic molecule with a solvent, characterizing the density of the amphiphile through a phase function *u* ∈ *H*^2^(Ω) via the free energy

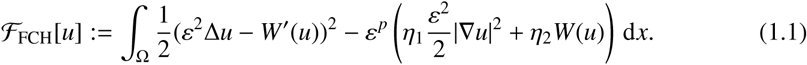

Here *W* is a double well potential with two unequal depth minima at *b*_−_ < *b*_+_ satisfying *W*(*b*_−_) = 0 > *W*(*b*_+_). The amphiphilic volume fraction is related to the density *u* − *b*_−_ with the equilibrium state *u* = *b*_−_ corresponding to pure solvent. The strength of the lower order functionalization terms are characterized by the value of *p*, generically selected as 1 or 2, and the values of *η*_1_ and *η*_2_. These parameters encode the affinity of the charged elements of the amphiphilic molecule for the solvent (called the solvent quality) and the aspect ratio of the amphiphilic molecule, respectively, see [2, 5, 9].

Experimental investigations show that when single-species amphiphilic materials are dispersed in solvent, called casting, they self-assemble into a diverse array of molecular-width structures, [10, 18]. The associated bifurcation diagram depends subtly upon both the aspect ratio of the amphiphilic molecule and the solvent quality. Molecules with aspect ratio near unity form two-molecule thick bilayer membranes familiar from cellular biology. Larger aspect ratio molecules form higher codimensional structures such as filaments and micelles and complex networks with triple junctions and end-caps. Within the casting experiments the genesis of this structural diversity has been referred to as the onset of ‘morphological complexity’, [17]. Gradient flows of the scalar FCH free energy provide an accurate representation of this bifurcation structure, providing a mechanism for the onset of morphological complexity via a transient passage through a pearling instability that leads bilayers to and break into filaments and other higher codimension morphologies, [9]. The single species bilayers supported by the scalar FCH free energy are always neutrally stable to pearling bifurcations at leading order – opening the door for lower order terms, including the system parameters *η*_1_ and *η*_2_ and the dynamic value of the bulk density of amphiphilic material to play a decisive role, [20]. Indeed, previous work on the scalar FCH has shown that the neutral modes of its bilayer interfaces are associated either to motion of the underlying interface, termed meander, or to the short wave-length modulations of their width associated to pearling, [16]. In regimes in which interfaces are stable to the pearling bifurcation, the interfacial motion has been rigorously described through a normal velocity proportional to curvature, with the proportionality constant depending upon the difference between the evolving bulk density of amphiphilic materials away from the interface and the bilayer bulk-density equilibrium value. Significantly this proportionality can be negative, which is typical in casting experiments in which the bulk density is high, and leads to a *curve lengthening* motion regularized by surface diffusion, [6].

In biologically relevant settings, phospholipid membranes are robustly stable to pearling bifurcations, which would generically be toxic to the living cell or to the organelle enclosed by the membrane. Significantly phospholipid membranes are never comprised of a single species. Generically significant amounts of cholesterol or other glycolipids are blended into the phospholipid membranes. Indeed all eukaryotic plasma membranes contain large amounts of cholesterol, often a 1-1 molar mixture of phospholipids and cholesterol [1]. While phospholipids are classic amphiphilic materials with a charged head group and a hydrophobic tail, cholesterol is a shorter, asymmetric molecule with a small, weakly charged head and a hydrophobic body. Within a phospholipid membrane cholesterol typically wedges itself in the void space between the amphiphilic phospholipid molecules, see Figure 1, where it significantly constrains the motion of the lipids.

**Figure 1:**
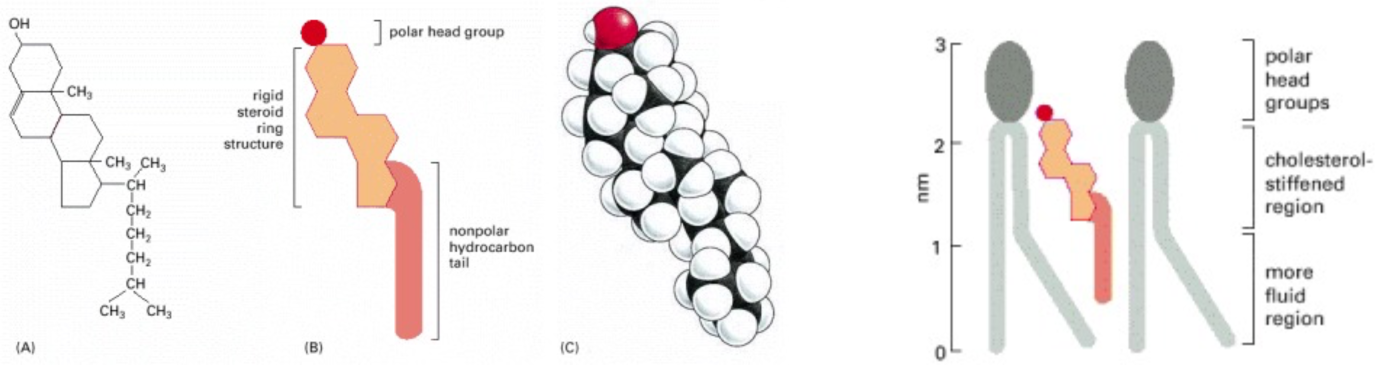
(left) (a) Chemical composition, (b) schematic, and (c) volume rendering of cholesterol. (Right) Caricature of cholesterol residing with void space of a lipid bilayer. Its small head group serves to orient the molecule and its asymmetric shape provides leverage on the lipid’s tail groups to constraint their range of motion. Reprinted with permission from [1]

We introduce the multicomponent functionalized energy as a general framework for a system of *n* + 1 constituent species residing in a domain Ω ⊂ ℝ^*d*^, and provide a sharp characterization of the bilayer structures that are robustly stable to pearling bifurcations. The characterization involves only the spectrum of the linearization of the reduced dynamical system (1.5) that defines the connecting profile. This framework contains the two-component singularly perturbed systems as a subfamily that describes strongly asymmetric two-component mixtures. Previous work has exploited the asymmetry to provide explicit leading order constructions of homoclinic connections [11]. In Theorems 3.5 and 3.9 we show that the robust pearling stability condition corresponds to a natural geometric feature arising in the singular perturbation construction. We develop a minimal two component phospholipid-cholesterol bilayer (PCB) model that mimics essential features of this ubiquitous system. In particular the PCB model encodes two strong asymmetries: the ratio of the lengths of cholesterol and phospholipid, and the lever-arm nature of cholesterol’s shape and its interdigitated packing that allows cholesterol to exert an outsized influence on the phospholipid tails [22]. We propose that these asymmetries afford the mechanism by which cholesterol type molecules robustly stabilize phospholipid membranes.

### 1.1 The Multicomponent Functionalized Energy

The multicomponent functionalized (MCF) energy takes the form

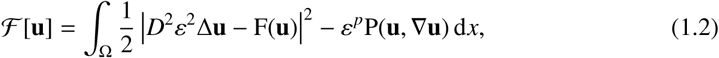

with **u** ∈ *H*^2^(Ω), *D* is an *n* × *n*, positive diagonal matrix, F : ℝ^*n*^ ↦ ℝ^*n*^ is a smooth vector field, and P : ℝ^*n*^×ℝ^*d*×*n*^ ↦ ℝ represents the lower order functionalization term. This model generalizes the multicomponent functionalized Cahn-Hilliard free energy introduced in [24], replacing the gradient form of the vector field with the more general function F whose non-gradient form plays a central role in the generation of robust pearling stability.

The components 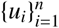 of **u** and *u*_*n*+1_ := 1−*u*_1_ −…−*u*_*n*_ represent the volume fractions of the *n* + 1 constituent species. Each species is classified as either amphiphilic or solvent. There can be more than one solvent phase, in which case they are generally immiscible, [4]. The critical points 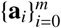 of F are associated to pure solvent phases and act as rest-states for the system. The domains Ω_*i*_ = {*x* ∈ Ω ||**u**(*x*) − **a**_*i*_| = *O*(*ε*)} can have *O*(1) volume without generating leading order contributions to the free energy. The dominant term in the multicomponent functionalized energy encodes proximity to “good packings” of the molecules identified as solutions, or approximate solutions, of the packing relation: *D*^2^*ε*^2^Δ**u** = F(**u**). The MCF energy is typically coupled with a non-negative linear operator 𝒢, called the gradient, that annihilates the constants. A canonical choice is 𝒢 = −Δ. The result is the gradient flow

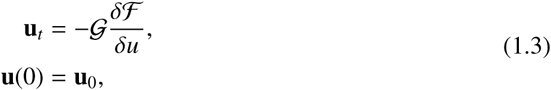

where the variational derivative is taken with respect to the *L*^2^(Ω) inner product. When combined with appropriate boundary conditions, for example periodic boundary conditions, the result is a flow which decreases the energy ℱ[**u**(*t*)] while preserving the total mass of each constituent species. This work focuses on the properties of the energy, and constructions that lead to normally coercive low-energy manifolds of ℱ.

In section 2 we characterize the properties of connecting orbits that arise as the good packings that separate domains Ω_*i*_ and Ω _*j*_ with an *O*(*ε*) width interface comprised of amphiphilic molecules. We take the interface to be flat, and measure normal distance in the scaled variable *z*(*x*) := dist(*x, ∂*Ω_*i*_)/*ε*, and drop the lower order functionalization term P, so that the connecting profiles can be characterized as minimizers of the codimension one reduced energy

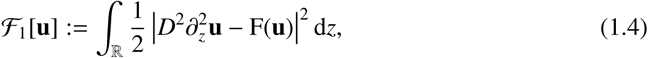

subject to the constraint **u** − *ϕ*_*i j*_ ∈ *H*^2^(ℝ) where *ϕ*_*i j*_ := **a** _*j*_ + (**a**_*i*_ − **a** _*j*_)(1 − tanh(*z*))/2 satisfies *ϕ*_*i j*_ → **a** _*j*_ as *z* → ∞ and *ϕ* → **a**_*i*_ as *z* → − ∞. When they exist, the absolute minimizers are the orbits of the 2*n* dimensional dynamical system

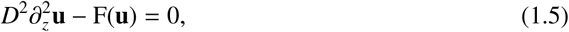

that are heteroclinic (or homoclinic) to the equilibria **a**_*i*_ and **a** _*j*_. These orbits are global minimizers of ℱ_1_, yielding zero energy. Correspondingly we call (1.5) the freeway system and the associated heteroclinic or homoclinic orbits the freeway connections. When the diagonal elements of *D* are strongly unequal, the freeway system fits within the framework of geometric singular perturbation (GSP) theory.

The local minimizers of the reduced free energy, for which the quadratic residual is not zero, also provide relevant connections between phases, especially when freeway connections do not exist. They satisfy

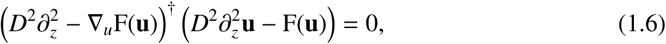

subject the heteroclinic or homoclinic boundary conditions. Here ^†^ denotes *L*^2^ adjoint, see subsection 1.2. This is a 4*n* dimensional dynamical system, and its solutions generically have non-zero reduced energy. We call this the toll-road system, and the associated heteroclinic orbits the toll-road connections. Even when matrix *D* has the singularly perturbed structure, the toll-road system does not trivially fit within the classical singularly perturbed framework. However we show that toll-road connections are generically generated at saddle-node bifurcations of freeway connections, and characterize the energy of the associated toll-road connection in terms of the saddle-node bifurcation parameter. These results establish the GSP theory as a powerful tool for the construction of MCF energies that support families of robustly stable connections with prescribed composition.

In section 3 we extend the zero-energy, flat-interface, freeway connections generated by the GSP theory to low-energy, curved-interface functions in *H*^2^(Ω) through a dressing process, given in Definition 3.2. This allows the construction of a low-energy, freeway manifold parameterized by underlying “admissible interfaces”, given in Definition 3.1. The main analytical result, Theorem 3.5, characterizes homoclinic freeway connections for which the associated freeway manifold is normally coercive, independent of *ε* sufficiently small. The principal loss of coercivity in scalar systems arises through the onset of the pearling bifurcation which triggers a high-frequency modulation of the bilayer width that can lead to its break-up into structures with lower codimension, [9]. Indeed, the pearling bifurcation can be triggered dynamically by *O*(*ε*) changes in the bulk lipid density. Theorem 3.5 specifically rules out these classes of instability through a condition on the spectrum of the linearization, L of the homoclinic freeway connection about the freeway system (1.5), see (3.10), that is readily verifiable within the GSP framework. There is a significant literature that develops rigorous estimates on slow motion of gradient flow systems near low-energy manifolds, see [23] and [3]. A key component of this analysis is played by the uniform coercivity of the energy to perturbations normal to the manifold, that allow the derivation of the asymptotic evolution of the system in the tangent plane of the manifold. In [14] these slow flow results have been extended to recover leading order dynamics associated to the slow flow, and we believe that the results of Theorem 3.5 will allow the interfacial motion results in [6] to be extended rigorously to a wide class of gradient flows of the MCF energy near the low-energy freeway manifolds constructed herein.

In Section 4 we examine the structure of the MCF energy in the neighborhood of a saddle node bifurcation of freeway homoclinics within the GSP framework. At the bifurcation point the kernel of L is not simple, rendering Theorem 3.5 inapplicable. Modulo non-degeneracy assumptions Theorem 4.2 shows that the freeway saddle node bifurcation induces a toll-road homoclinic and characterizes its energy as quadratic function of the distance of the bifurcation parameter past criticality. In particular, we give an explicit example of a freeway saddle node bifurcation within the PCB model, characterizing the energy of the toll-road homoclinic in terms of the readily computable geometric features of the model.

The synergy between the MCF energy and the GSP theory is particularly fortuitous, as there is limited intuition for the relation between the structure of the nonlinearity in higher-order, multicomponent models and the physical properties of the constituent molecules. Rigorous derivation of higher-order free energies from more fundamental models, such as the derivation of the Ohta-Kawasaki free energy from the self-consistent mean field theory, have been performed, see [7, 8] for a general framework and [27, 28] for models specific to surfactants. However the analysis in such derivations is generically weakly nonlinear, and affords little information on nonlinear interactions beyond those imposed in an ad-hoc manner, generically through incompressibility arguments. The MCF energies constructed from the GSP approach are strongly nonlinear and strongly asymmetric in their nonlinear terms. This asymmetry plays an essential role in the analysis, rendering the operator L strongly non-selfadjoint and sweeping its spectrum off of the positive real axis and into the complex plane. This complexification is stabilizing as neutral modes in the linearization of the MCH about a homoclinic freeway connection arise from a balance between positive real spectrum of L against negative spectrum of the surface diffusion operator.

The phospholipid-cholesterol bilayer model presented in Section 2.3, is the minimal GSP based model that supports both a single-phase pearling-neutral phospholipid bilayer, and a two-phase phospholipid-cholesterol bilayer that is robustly stable to pearling. It is tempting to find synergy between the generic, geometric nature of these stability results and the generic presence of cholesterol within phospholipid membranes. Cholesterol’s interdigitation between lipid molecules leads to a core density peak and an outsized impact on lipid mobility that inhibits the lipid tail compression required for the onset of pearling bifurcations, [22]. It may be that the genome has latched upon the generic, geometric, singular role of cholesterol as a mechanism to prevent formation of micelles and other higher codimensional defects within phospholipid membranes.

### 1.2 Notation

Consider a function *f* : ℝ ↦ *X* where *X* is a Banach space and *s* ∈ ℝ is a parameter in *f*. We say that *f* is *s*-exponentially small in *X* if there exists *ν* > 0 such that ‖*f*‖_*X*_ ≤ *e*^−*ν*/*s*^ for *s* > 0 as *s* ≪ 1 tends to zero.

We use ^*t*^ to denote the transpose of a matrix or a vector in the usual Euclidean inner product and ^†^ to denote the an adjoint operator or eigenfunction in the *L*^2^(Ω) inner product.

## 2 Connecting Orbits

In this section we establish the structure of the freeway and toll-road connection problems and the existence of specific solutions in the context of the geometric singular perturbation scaling.

### 2.1 Freeway and Toll-road connections

We assume that F : ℝ^*n*^ ↦ ℝ^*n*^ is smooth and has *m* + 1 critical points **a**_0_, …, **a**_*m*_ for which F(**a**_*i*_) = 0, and *D* is an *n* × *n*, non-negative diagonal matrix. Generically the phase space is mapped onto species densities with the variable *u*_*i*_ denoting the volume fraction of species *i*, residing in

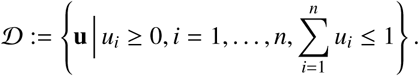

Critical points of *F* denote the solvent phases, and when modeling a mixture with a single solvent it is generically taken as **a**_0_ := (0, …, 0) with {*u*_1_, …, *u*_*n*_} denoting *n* amphiphilic phases. In low energy configurations these surfactants reside on thin interfaces generically of codimension one or higher, that are *O*(*ε*) thin in one or more directions (the co-dimensions). We focus on codimension one geometries, and in this section fix the interface Γ to be a flat *d* − 1 dimensional hypersurface, so that the minimization problem reduces at leading order to the system for ℱ_1_ given in (1.4). The infimum is non-negative and if attained, then the minimizer is smooth and satisfies the associated Euler-Lagrange equation (1.6), which we call the toll-road system. Setting G = *D*^−2^F, it is convenient to write the toll-road system as a 4*n* dimensional, first-order system,

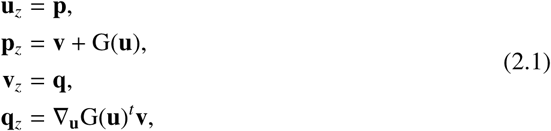

where as a consequence **v** := **u**_*zz*_ − G(**u**)^*t*^. An equilibrium **a** of *F* is normally hyperbolic if the linearization about the equilibrium **A** := (**a**, 0, 0, 0)^*t*^ ∈ ℝ^4*n*^ of (2.1) has no purely imaginary eigenvalues.

#### Lemma 2.1.

*An equilibrium* **a** ∈ ℝ^*n*^ *of F is normally hyperbolic (2.1) if and only if the n* × *n matrix D*^−2^ ∇_**u**_ *F*(**a**) *has no eigenvalues in the set* ℝ_−_ = (−∞, 0]. *In this case* **A** = (**a**, 0, 0, 0)^*t*^ *has a* 2*n dimensional stable and* 2*n dimensional unstable manifold within (2.1). The system (2.1) has a conserved quantity* H : ℝ^4*n*^ ↦ ℝ *given by*

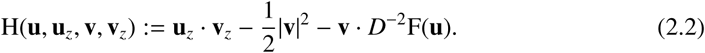

*In particular homoclinic and heteroclinic solutions of (2.1) lie on the* ℝ^4*n*−1^ *dimensional* {*H* = 0} *level set. Let* **a**_*i*_ *and* **a** _*j*_ *be two normally hyperbolic equilibria and* Φ = Φ_*i j*_(*z*; *γ*) *be a smooth k* ≥ 1 *dimensional manifold of connections between* **a**_*i*_ *and* **a** _*j*_. *Then there exists α*_*i j*_ ∈ ℝ_+_ *such that* ℱ_1_(Φ) = *α*_*i j*_ *for all connecting orbits* Φ *on the manifold.*

*Proof.* Using the relation **v** = **u**_*zz*_ − G(**u**)^*t*^ we take the dot product with **u**_*z*_

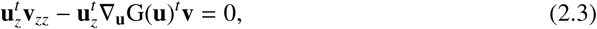

and equivalently, since a scalar equals its own transpose, we have

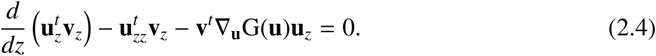

Substituting **u**_*zz*_ = **v** + G(**u**) we find

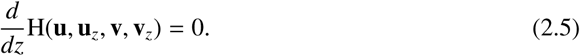

where H is as defined in (2.2).

Each of the critical points **a** of F satisfies H(**a**, 0, 0, 0) = 0, and since H is conserved under the flow the orbits connecting these critical points reside on the 4*n* − 1 dimensional level set {H = 0}. If **a** is a critical point of G then the linearization of the system (2.1) about **A** := (**a**, 0, 0, 0)^*t*^, takes the form **U**_*z*_ = M**U**, where

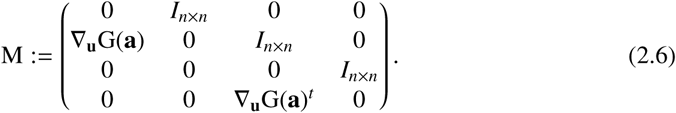

We compute that *λ* ∈ *σ*(**∇**_**u**_(G(**a**))) if and only if the four values 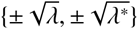 lie in *σ*(M) up to algebraic multiplicity. In particular the isolated critical point **a** of F is normally hyperbolic within the toll-road system if and only if *D*^−2^ **∇**_**u**_(F(**a**)) has no purely real, negative eigenvalues. If **a** is normally hyperbolic, the symmetry of the spectrum of M guarantees that the stable and unstable manifolds of the 4*n* dimensional system has equal dimension, hence they are both 2*n* dimensional.

To establish the uniformity of the energy over the manifold of connections, we insert Φ_*i j*_ into (1.6) are rewrite it in the form

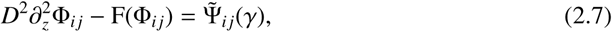

where the right-hand side, 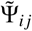, lies in the kernel of the adjoint of

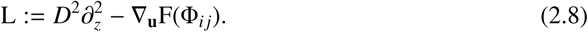

Taking the partial of (2.7) with respect to *γ* yields the relation

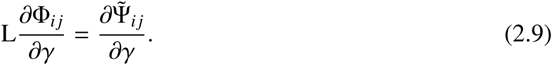

This implies that the right-hand side is *L*^2^(ℝ) orthogonal to the kernel of L^*t*^ and hence to 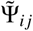. In particular 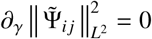 and we may write 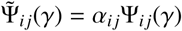 where 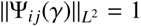. The result follows since 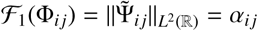 is independent of *γ*. □

We reinforce this dichotomy of zero-energy and non-zero energy connections through the following definition.

#### Definition 2.2.

*If a manifold of connections* Φ_*i j*_ *has zero energy, α*_*ij*_ = 0, *then the constituent orbits satisfy the freeway sub-system (1.5). We call these orbits freeway connections.*

### 2.2 Freeway homoclinic connections in singularly perturbed systems

Establishing the existence of connections in *n*-dimensional dynamical systems of the general form (1.5) is nontrivial. However, when the eigenvalues of the matrix *D* exhibit a wide range of magnitudes, controlled by a small parameter 0 < *δ* ≪ 1, then the associated dynamical system may have orbits that can be rigorously constructed via geometric singular perturbation theory by gluing together solutions of the so-called slow and fast sub-systems of reduced dimension. In [11], theory was developed that provides for the existence and spectral analysis of homoclinic connections in a general class of two-component, singularly perturbed vector fields for the case in which the vector field is strongly non-symmetric. The homoclinic connection problem is equivalent to the freeway system (1.5) with *n* = 2, and *D* = diag(1, *δ*), where 0 < *δ* ≪ 1, and the vector field F takes the form

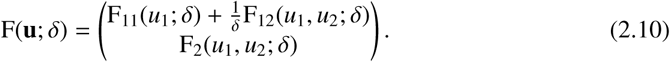

The component functions F_*i j*_ obey mild regularity assumptions [11]. The resulting model can be written as a first order dynamical system in the form (1.6), which in the fast spatial variable *ζ* := *z*/*δ* takes the form

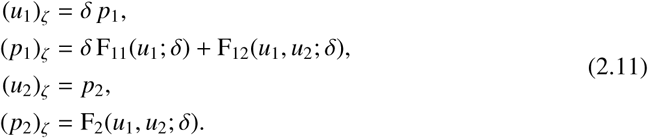

We require that the following structural assumptions hold:

#### Assumption 2.3.

*The point* **a** = (0, 0) *is an isolated, hyperbolic equilibrium of* (1.5). *The component functions satisfy* F_12_(*u*_1_, 0; *δ*) = 0 *and* F_2_(*u*_1_, 0; *δ*) = 0 *for every u*_1_ ∈ ℝ. *There exists an open set V* ⊂ ℝ, *such that the planar system* (*u*_2_)_*ζζ*_ − F_2_(*s, u*_2_; 0) = 0 *admits a symmetric solution u*_2,*h*_(*ζ*; *s*) *that is homoclinic to u*_2_ = 0 *for every s* ∈ *V.*

#### *Remark* 2.4.

The parameter *δ* is taken asymptotically small in the GSP theory, however in the context of the MCF energy it denotes the ratio of lengths comparable molecules, and is not vanishingly small. Correspondingly we take *δ* sufficiently small to apply the GSP theory, but then consider it to be a fixed parameter in the subsequent analysis of the MCF energy. In particular the upper bound *ε*_0_, on the value of admissible *ε* will depend upon the fixed value of *δ*. In effect the GSP theory applies in the regime *ε* ≪ *δ* ≪ 1, as is consistent with applications.

From these assumptions, it directly follows that the origin of (2.11) is a hyperbolic equilibrium. Moreover, we see that the manifold ℳ_0_ := {*u*_2_ = *p*_2_ = 0} is invariant under the flow of (2.11). The flow on ℳ_0_, which we call the reduced slow flow, is given to leading order in *δ* in the slow variables by

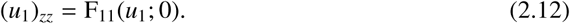

From the assumptions, the point (*u*_1_, *p*_1_) = (0, 0) is a hyperbolic equilibrium of (2.12) and the associated (slow) stable and unstable manifolds 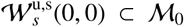 are one-dimensional, and equal to the other’s reflection about the *u*_1_-axis; see Figure 2. Defining 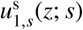 as the unique positive solution to (2.12) satisfying 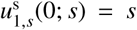 and 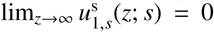, we see that the one-dimensional (slow) stable manifold of the origin 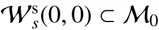 for *u*_1_ > 0 is given by the orbit of 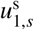. Moreover, both the stable and unstable manifolds, 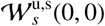, lie on the level set 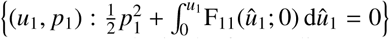.

**Figure 2:**
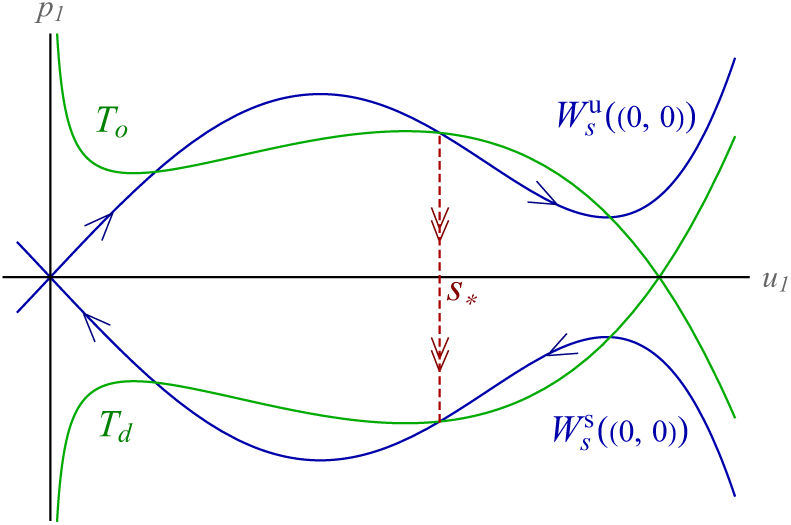
A schematic representation of the reduced slow flow on ℳ_0_. The slow stable and unstable manifolds 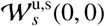 are indicated in blue, the take-off and touchdown curves *T*_o_ and *T*_d_ are indicated in green. The jump through the fast field at a transversal intersection of 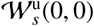 and *T*_o_ for *u*_1_ = *s*_*_ is indicated in red.

Conversely, in the fast scaling (2.11), we see that to leading order in *δ, u*_1_ = *s* is constant,

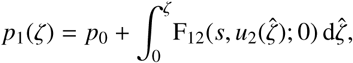

while *u*_2_ obeys the so-called reduced fast flow

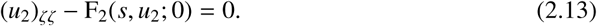

The manifold ℳ_0_ is exactly the set of trivial equilibria of (2.13); by the requirements of Assumption 2.3, these trivial equilibria are hyperbolic. Moreover, there exists an open subset ℳ_1_ ⊂ ℳ_0_,ℳ_1_ := {*u*_2_ = *p*_2_ = 0, *u*_1_ ∈ *V*}, such that the reduced fast flow connects (*s, p*_1,*o*_, 0, 0) _1_∈ ℳ to (*s, p*_1,*d*_, 0, 0) ∈ ℳ_1_ through the symmetric fast homoclinic orbit *u*_2,*h*_(*ζ*; *s*). In this reduced limit, the jump in *u*_1_’s derivative satisfies

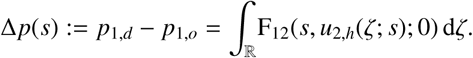

This suggests the definition of a pair of curves on ℳ_0_, called the “take-off” and “touchdown” curves, for which Δ*p* transports the orbit first away from, and then back to ℳ_0_ in a symmetric fashion. The take-off curve is given by 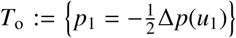, while the touchdown curve *T*_*d*_ is given by its reflection about the *u*_1_-axis; see Figure 2.

A homoclinic orbit of the GSP scaling of the 4-dimensional freeway problem lies in the transversal (first) intersection of the 2-dimensional stable and unstable manifolds of the origin (0, 0, 0, 0). The scale separation present in the system allows us to decompose this intersection into a first slow segment that follows 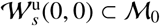 closely, then makes a fast excursion away from ℳ_0_, but 𝒪(*δ*) close to *u*_2,*h*_(*ζ, s*_*_) for some *s*_*_, and then touches down again near ℳ_0_ to closely follow 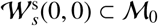 back to the origin (0, 0, 0, 0). In the singular limit, this concatenation procedure provides a homoclinic orbit precisely when the take-off curve *T*_o_ intersects the slow unstable manifold 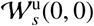; see Figure 2. When this intersection is nongenerate, transversality arguments imply that the singular orbit persists for sufficiently small 0 < *δ* ≪ 1; for the full analysis, see [11].

We define the function *ρ* : *V* ↦ ℝ

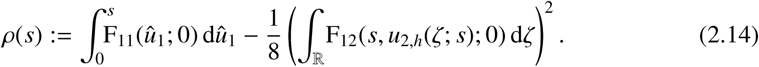

One can deduce that if the take-off curve *T*_o_ and the slow unstable manifold 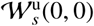 intersect transversally at *u*_1_ = *s*_*_, then the function *ρ* has a nondegenerate root at *s* = *s*_*_. However, *ρ* also vanishes when *T*_o_ intersects the slow *stable* manifold 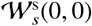, which does not lead to a meaningful geometric construction when 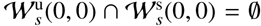. To exclude these spurious roots, we employ the explicit characterisation of 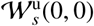 by the solution 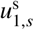, and introduce the condition that 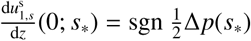.

The following is a reformulation of [11, Theorem 2.1].

#### Theorem 2.5.

*Assume n* = 2, *D* = diag(1, *δ*), *that* F *takes the form* (2.10), *and that the cond tions of Assumption 2.3 hold. Fix δ* > 0 *sufficiently small. Let N denote the number of nondegenerate roots of ρ, defined in* (2.14), *that lie in the set V, and that obey the condition*

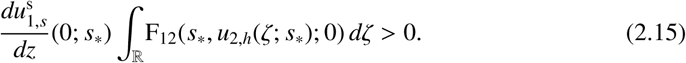

*Here*, 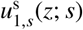 *is the unique positive solution to* (2.12) *satisfying* 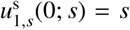 *and* 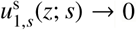. *as z* → ∞ *Then there are N symmetric, positive, one-circuit solutions to* (1.5) *that are homoclinic to* **a** = (0, 0). *In particular, for each root s*_*_ *the associated homoclinic connection* (*u*_1,*_(*z*), *u*_2,*_(*z*)), *translated to be even about z* = 0, *has the following spatial structure:*

1. *for* 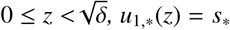 *and u*_2,*_(*z*) = *u*_2,*h*_(*z*/*δ*; *s*_*_) *to leading order in δ;*
2. *for* 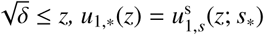 *to leading order in δ, while u*_2,*_(*z*) *is δ-exponentially small.*

#### *Remark* 2.6.

The result from [11] encompasses a larger class of systems, in particular the equilibrium **a** may lie on the boundary of the domain of definition of the vector field F. This necessitates additional technical assumptions on F, see [11, Assumptions (A1-4)].

### 2.3 A minimal phospholipid-cholesterol model

We apply the singularly perturbed framework presented in section 2.2 to develop a minimal model of a phospholipid-cholesterol bilayer (PCB) membrane that supports both a pearling neutral bilayer membrane in the absence of cholesterol and a robustly stable membrane with an optimal phospholipid-cholesterol balance. The minimal PCB model takes the form

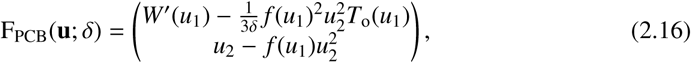

where *f* is a smooth, positive, non-increasing function. The slow component, *u*_1_, denotes the volume fraction of phospholipid while the fast component, *u*_2_, denotes that of cholesterol. The take-off curve *T*_o_ –which is directly present in the model formulation– is smooth and specified below. The scalar potential *W* is precisely the smooth double-well from (1.1) with minima at *b*_−_ = 0 and *b*_+_ = 1 satisfying *W*(0) = 0 > *W*(1). In particular *W*′(*u*_1_) has a unique transverse zero *u*_1_ = *u*_1,max_ that lies in (0, 1), so that the slow stable and unstable manifolds 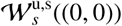 coincide for *u*_1_ > 0, leading to the existence of a fully slow homoclinic orbit on ℳ_0_. Moreover, since *f* is positive everywhere, the fast subsystem (2.13) admits a homoclinic orbit for every value of *u*_1_ = *s*, hence we may take *V* = ℝ in Assumption 2.3.

The homoclinic orbit *u*_2,*h*_ in the fast system (2.13) has the explicit expression

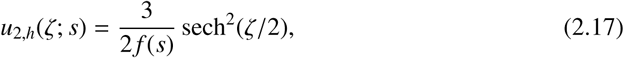

allowing us to evaluate the integral

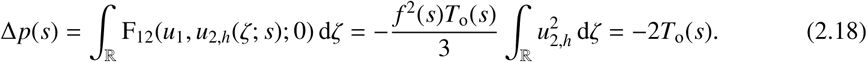

Since F_11_(*u*_1_; 0) = *W*′(*u*_1_) and *W*(0) = 0 the first integral in (2.14) takes the values *W*(*s*). Moreover, the portion of the unstable slow manifold in the first quadrant can be given as the graph {(*s*, ω_*u*_(*s*)) | *s* ∈ (0, *u*_1_)}, where 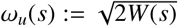. We calculate that

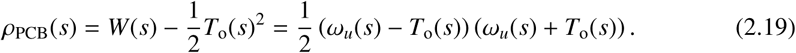

As established in section 2.2, the zeros of *ρ*_PCB_ correspond to the crossings of the take-off curve with the graph of the unstable slow manifold. We choose the take-off curve to have a transverse intersection with the unstable manifold at the phospholipid density *u*_1_ = *s*_*_ corresponding to a bilayer membrane fully interdigitated with cholesterol. The cholesterol density is modulated by adjusting the value of *f* (*s*_*_), see (2.17). For the slow subsystem, the slow stable and unstable manifolds of the origin 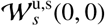 coincide, so that condition (2.15) is automatically satisfied – that is, *every* root of *ρ*(*s*) is a valid candidate for the construction outlined in section 2.2. In Figure 3, the dynamics on ℳ_0_ and a corresponding pulse are shown for a specific choice of *T*_o_.

**Figure 3:**
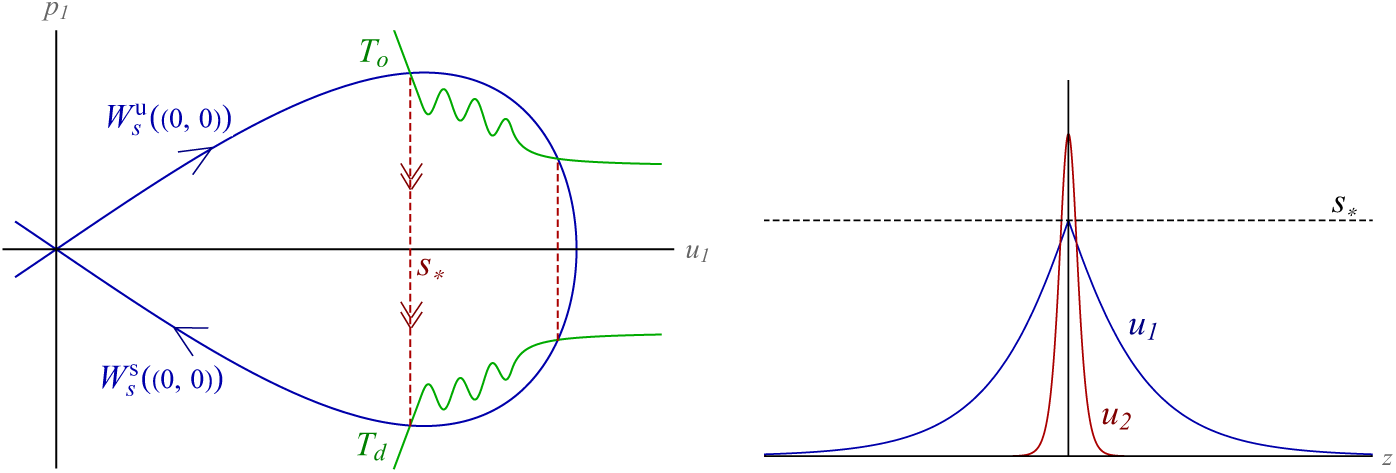
(left) The phospholipid cholesterol bilayer system (2.16) has two fast-slow homoclinic freeway connections, corresponding to the two intersections of the take-off curve *T*_o_ with the unstable manifold 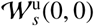 of the origin of the slow system (*u*_1_)_*zz*_ = *W*′(*u*_1_) (2.12). The left-most intersection, at *u*_1_ = *s*_*_ has *ρ*′(*s*_*_) > 0 and will yield robustly stable bilayer interfaces. The rightmost intersection yields a fast slow connection with *ρ*′ < 0. The system also supports a slow-only homoclinic with *u*_2_ = 0. (right) Depiction of the fast-slow homoclinic connection for the *u*_1_ = *s*_*_ intersection, corresponding to a cross section of a bilayer membrane with phospholipid (*u*_1_) on the outside and interdigitated cholesterol (*u*_2_) in the core. The maximum value of the slow component occurs at the take-off intersection point. The fast component *u*_2_ = *u*_2,*h*_(*ζ*) is scaled by *f* (*s*_*_).

## 3 Normal Coercivity of Homoclinic Freeway Manifolds

In this section we extend the freeway connections, generating the freeway manifold of low energy solutions associated to a wide class of admissible interfaces. We identify conditions which guarantee the normal coercivity of homoclinic freeway manifolds, and relate the stability conditions back to the construction of the homoclinic freeway connections within the GSP context.

### 3.1 Freeway Manifolds

We consider a codimension one interface Γ given by the local parameterization

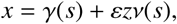

with *γ* : **S** ⊂ ℝ^*d*−1^ ↦ Γ ⊂ Ω and *ν*(*s*) the outward normal to Γ at *γ*(*s*). The pair (*s, z*) forms a local coordinate system, for which *s* = *s*(*x*) parameterizes location on Γ and *z* = *z*(*x*) is the *ε*-scaled signed distance to Γ. The line segments {*γ*(*s*) + *tν*(*s*) | |*t*| < *ℓ*}_*s*∈**S**_ are called the whiskers of length *ℓ* of Γ.

#### Definition 3.1.

*For any K, l* > 0 *the family* 𝒢_*K,l*_ *of admissible interfaces is comprised of closed (compact and without boundary), oriented d* − 1 *dimensional manifolds* Γ *embedded in* Ω ⊂ ℝ^*d*^, *which are far from self-intersection and posses a smooth second fundamental form. More precisely*, 2*ℓK* < 1, *the W*^4,∞^(**S**) *norm of the principal curvatures of* Γ *is bounded by K, the whiskers of length* 2*ℓ do not intersect each-other nor the boundary of* Ω, *and the surface area*, |Γ|, *of* Γ *is bounded by K. We call the set* Γ_2*ℓ*_ := {*x* ||*z*(*x*)| < 2*ℓ*/*ε*}, *the reach of* Γ.

For an admissible Γ the change of variables *x* ↦ (*s, z*) is a *C*^4^ diffeomorphism of Γ_2*ℓ*_, see section 6 of [16]. To each class 𝒢_*K,ℓ*_ we associate a symmetric, compactly supported 𝒞^∞^ function *ξ* that is monotone on ℝ_+_ and takes values 1 on [−*ℓ, ℓ*] and is 0 on ℝ\[−2*ℓ*, 2*ℓ*].

#### Definition 3.2.

*For each function* **u** *which tends at an exponential rate to constant values* **u** ↦ **u**_±_ *as z* ↦ ±∞, *we define its* Γ*-dressing as the L*^2^(Ω) *function*

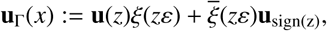

*where* 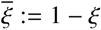.

We will denote both the Γ-dressing and the original *L*^2^(ℝ) function by **u** where doing so does not introduce confusion. The function *ξ*(*εz*(*x*)) lies in *H*^4^(Ω), even though the distance function *z* is not smooth outside the set Γ_2*ℓ*_.

#### Definition 3.3.

*To each freeway connection* **u**_*_ *of the subsystem (1.5) and admissible family of interfaces* 𝒢_*K,ℓ*_ *we associate the corresponding freeway manifold*

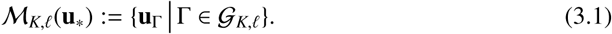

*comprised of the dressings of the admissible interfaces by* **u**_*_.

On the reach of Γ the (*s, z*) coordinate system induces a representation of the Cartesian Laplacian (denoted Δ_*x*_ to avoid ambiguity) in the form

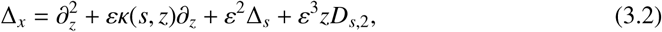

where κ(*s, z*) = *H*(*s*) + *O*(*εz*) is an extension of the mean curvature *H*(*s*) of Γ, Δ_*s*_ is the Laplace-Beltrami operator associated to Γ, and *D*_*s*,2_ is a second order operator in ∇_*s*_ with coefficients whose *W*^4,∞^ norm is bounded by *K*, see Proposition 6.6 of [16]. The eigenfunctions of Δ_*s*_ are given by 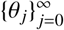 with eigenvalues 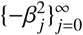 which satisfy 0 = *β*_0_ < *β*_1_ ≤ *β*_2_ …, see [21].

For functions supported in Γ_2*ℓ*_, the *L*^2^ inner product takes the inner form

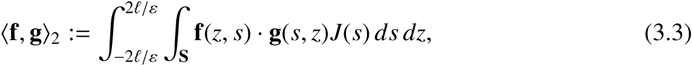

where *J* is the Jacobian of the change of coordinate map from *x* to (*z, s*). In particular 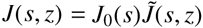 where *J*_0_ is the square root of the determinant of the first fundamental form of Γ and 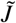 admits the expansion

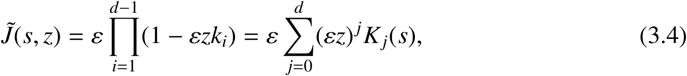

where *k*_1_, …, *k*_*d*−1_ are the principal curvatures of Γ, while *K*_0_ = 1 and for *i* = 1, … *d* − 1, *K*_*i*_ = *K*_*i*_(*s*) are (−1)^*i*^ times the sum of the all products of *i* curvatures of Γ. From the condition 2*ℓK* < 1 for interfacial admissibility we deduce that

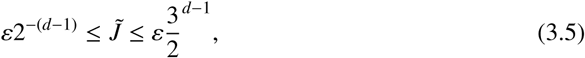

and hence after a scaling by *ε*, the inner product

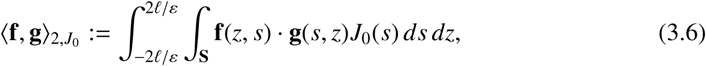

induces a norm equivalent to the usual *L*^2^ norm on Γ_2*ℓ*_. The Laplace-Beltrami eigenmodes are orthonormal in the inner product

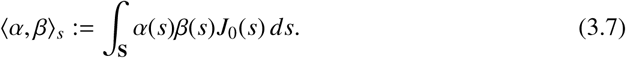

#### Proposition 3.4.

*Let* **u**_*_ *be a freeway connection between two equilibrium* **a**_−_ *and* **a**_+_ *of* F. *The freeway manifold associated to* **u**_*_ *lies in H*^2^(Ω). *If the functionalization term within the multicomponent functionalized energy (1.2) satisfies* P(**a**_±_, 0) = 0, *then*

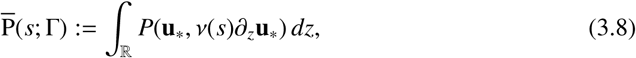

*is finite, and the dressings* **u** = **u**_Γ_ ∈ ℳ_*K,ℓ*_(**u**_*_) *have leading order energy*

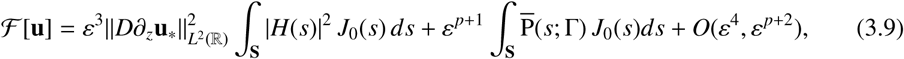

*where H denotes the mean curvature and ν the outer normal of* Γ.

The proof of this result is a direct modification of prior results, see [12, eqn (3.3) and Proposition 4.1], and is omitted.

### 3.2 Normal coercivity

In the sequel we assume that **a**_0_ = 0 is a hyperbolic equilibrium of the vector field F, and that **u**_*_ is a freeway connection homoclinic to **a**_0_. Without loss of generality we may set the functionalization term P equal to zero as it lower order in *ε* and does not impact the *ε*-uniform coercivity bounds we seek in (3.18), see also Remark 3.6. We let L denote the linearization of the freeway system (1.5) about **u**_*_,

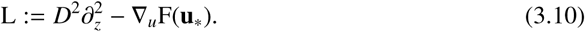

Let **u**_*_ = **u**_*_(Γ) denote the dressing of an admissible interface Γ by the homoclinic freeway connection **u**_*_. Our goal is to derive estimates on lower bounds of the spectrum of the second variational derivative of ℱ at **u**_*_, given by the operator

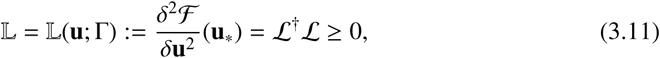

where we have introduced

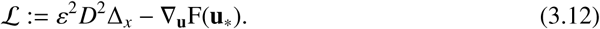

Within the reach of Γ the operator admits the exact expansion

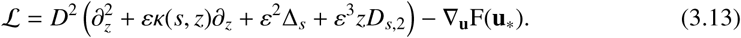

This motivates the introduction of

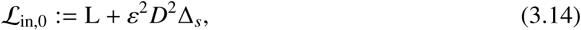

and the inner decomposition of ℒ as

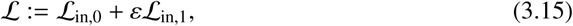

where ℒ_in,1_ := *D*^2^(κ(*s, z*)*∂*_*z*_ + *ε*^2^*D*_*s*,2_). On the complement of the reach of Γ the operator ℒ is an *ε*-exponentially small perturbation of the *constant-coefficient* operator

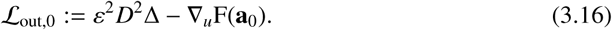

While 𝕃 from (3.11) is self-adjoint, its factor ℒ is not, indeed the spectrum of 𝕃 is precisely the singular values of ℒ. The spectrum of 𝕃 is clearly real and non-negative. We define the space of meander modes which closely approximate an *N*-dimensional subspace of the tangent plane of ℳ_*K,ℓ*_(**u**_*_),

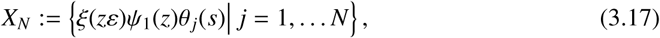

where the translational eigenfunction *Ψ*_1_ = *∂*_*z*_*u*_*_ ∈ ker(L). We also introduce 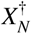 which is the adjoint space obtained by replacing *Ψ*_1_ with the adjoint eigenfunction 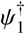. We show that for *N* sufficiently large that 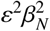 is *O*(1), then the operator 𝕃 is coercive on 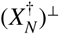, uniformly in *ε*, if the spectrum of *D*^−2^ *L* as an operator on *L*^2^(ℝ) has no strictly positive real spectrum.

#### Theorem 3.5.

*Let* **a**_0_ = 0 *be a normally hyperbolic equilibrium of* F *and let* **u**_*_ *be a solution of the freeway system (1.5) which is homoclinic to* **a**_0_, *and let* ℳ_*K,ℓ*_ *be the associated freeway manifold. Let the operators* L = L(**u**_*_) *and* 𝕃 = 𝕃(**u**_*_; Γ) *be as given in (3.10) and (3.11) respectively. If σ*_*p*_(*D*^−2^L) ∩ℝ_+_ = {0} *with a simple kernel spanned by ∂*_*z*_**u**_*_, *then for ℓK sufficiently small and for any fixed γ*_0_ > 0, *there exists ε*_0_, *μ* > 0 *such that for all ε* ∈ (0, *ε*_0_) *and all* Γ ∈ 𝒢_*K,ℓ*_

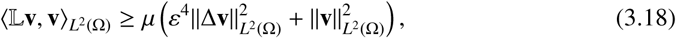

*for all* 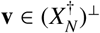 *where N* = *N*(*ε*) *is chosen to satisfy* 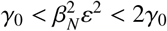.

#### *Remark* 3.6.

The coercivity extends to any *O*(*ε*) regular perturbation of 𝕃. Specifically the functionalization terms *ε*^*p*^P in ℱ add an *O*(*ε*^*p*^) regular perturbation to 𝕃 that does not impact the coercivity. Neither the functionalization terms nor perturbations to the form of **u**_*_ will impact the coercivity, and specifically they cannot induce the pearling bifurcations to which the freeway manifolds of the scalar FCH are susceptible.

*Proof.* The operator 𝕃 admits distinct formulations when acting on functions supported in Γ_2*ℓ*_ and on those supported in Ω \ Γ_2*ℓ*_. We decompose 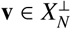 as

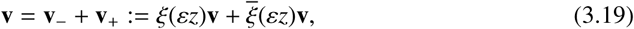

and writing 𝕃 = ℒ^†^ℒ, we expand the left-hand side of (3.18) as

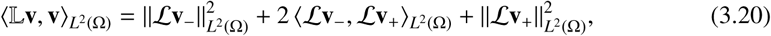

denoting the summands on the right-hand side as the inner, mixed, and outer bilinear terms respectively, we estimate them individually.

#### Inner bilinear term

From the inner formulation, (3.15), of 𝕃 we focus on the leader-order operator 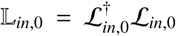, and consider it as acting on functions defined on the abstract set **S**_∞_ := **S** × ℝ formed from the unbounded whiskers. This is not a subset of Ω as the whiskers of length greater than 2*ℓ* generically intersect. We first establish the coercivity of the operator 𝕃_*in*,0_ in the *L*^2^(**S**_∞_) norm, defined as

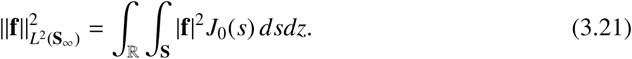

As observed in (3.5), for admissible interfaces Γ ∈ 𝒢_*K,ℓ*_ there exists *c* > 0 such that

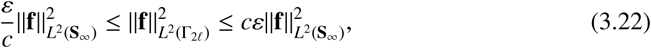

for all **f** ∈ *L*^2^(Γ_2*ℓ*_). We also introduce function space *H*^2^(**S**_∞_) with norm given by

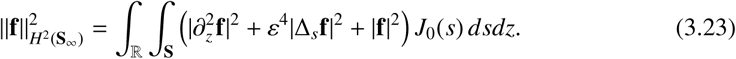

For functions supported in Γ_2*ℓ*_ the inner expression for *ε*^2^Δ given in (3.2) affords the estimates

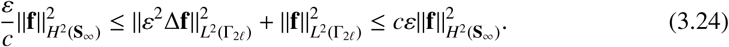

##### Lemma 3.7.

*There exist μ*_0_ > 0 *such that for all ε* ∈ (0, *ε*_0_) *and all* Γ ∈ 𝒢_*K,ℓ*_

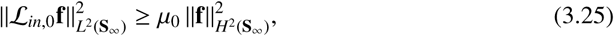

*for all* 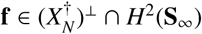

*Proof.* We decompose

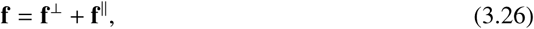

where **f**^‖^the component of **f** that lies in the *L*^2^(**S**_∞_)-tangent plane to *X*_*N*_, precisely,

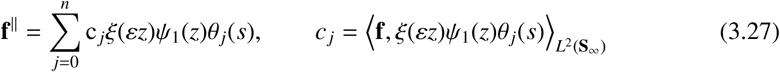

*Step 1: Control of* **f**^‖^*in H*^2^(**S**_∞_). Since the 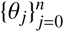 are orthonormal in *L*^2^(**S**) and 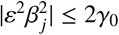 for all *j* = 0, …, *n*, we have the estimate

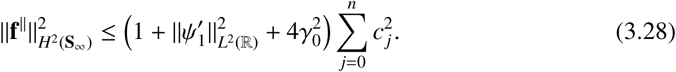

From the assumption **f**⊥*ξΨ*θ _*j*_ in *L*^2^(Ω) and the inner product representation (3.3), we derive

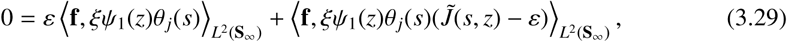

which allows us to rewrite the expression for *c* _*j*_ in (3.27) as

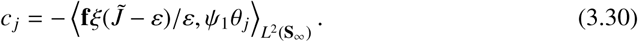

We carry out the *z* integral θ _*j*_ and express the remainder in the inner product (3.7),

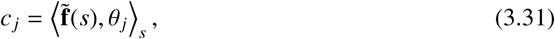

where we have introduced

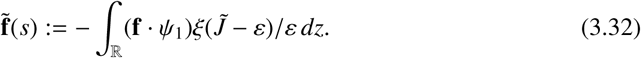

Using Hölder’s inequality we bound

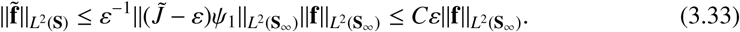

For the last estimate we employed the second identify in (3.4) together with the fact that the higher order curvatures *K*_*j*_ are uniformly bounded and and *Ψ*_1_ converges to zero exponentially in |*z*| so that |*z*|^*k*^*Ψ*_1_ is uniformly bounded in *L*^2^(ℝ) for *k* = 0, …, *d* −1. The functions {θ _*j*_(*s*)} are orthonormal in the inner product (3.7), so Plancherel’s identity implies

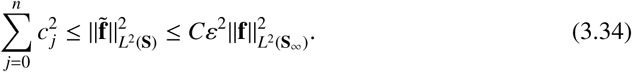

for a different constant *C*. In particular, from (3.22) and (3.28) it follows that

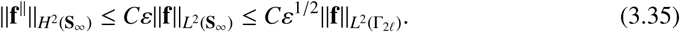

*Step 2: L*^2^(**S**_∞_)*-coercivity of* ℒ_in,0_ *on* 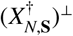. The spaces *X*_*N*,**S**_ and 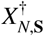 are the analogues of *X*_*N*_ and 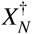 in *L*^2^(**S**_∞_) obtained by dropping the *ξ* cut-off in (3.17). We establish the coercivity

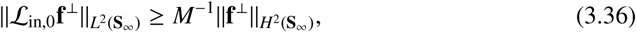

for 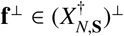 through the equivalent estimate

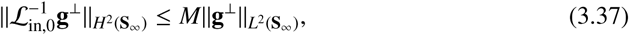

for all 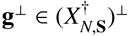. Since 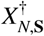 is comprised of eigenspaces of 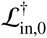 it follows that the condition 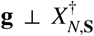 follows if and only if 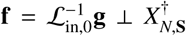. In the remainder of step 2 we drop the ⊥ superscript on **f** and **g**, bounding **f** ∈ *H*^2^(**S**_∞_) where **f** solves

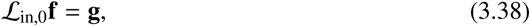

subject to 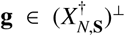. We decompose **g** and **f** into their inner Fourier components via the decomposition

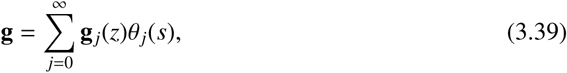

where the inner coefficients are given via formula

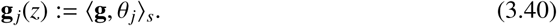

using the inner product from 3.7. This yields the uncoupled sub-problems

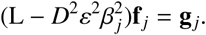

Since *u*_*_ is a homoclinic and defined on all ℝ, the operators have natural extension to *L*^2^(ℝ). We replace 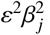 with *k* and define the family of operators {L_*k*_}_*k*≥0_, where L _*k*_ := (L − *D*^2^*k*). For each **h** ∈ *L*^2^(ℝ) we form the function

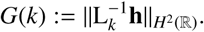

By the spectral assumption of Theorem 3.5, the operator L = L_0_ has a simple eigenvalue at 0, which is removed by the projection off of 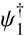, while for all *k* > 0, L_*k*_ is invertible from *L*^2^(ℝ) in *H*^2^(ℝ). For *j* = 0, …, *N*(*ε*), corresponding to 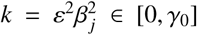, we have 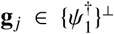. Consequently, for *k* ∈ [0, *γ*_0_] we consider 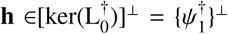 so that *G* is defined, finite at *k* = 0 and continuous in *k* on [0, *γ*_0_]. Since [0, *γ*_0_] is compact, *G* is uniformly bounded on this set for each **h**. From the uniform boundedness principal we conclude that the operators 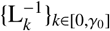 are uniformly norm bounded from {*Ψ*_1_}^⊥^ ⊂ *L*^2^(ℝ) to *H*^2^(ℝ). For *k* ≥ *γ*_0_ we consider **h** ∈ *L*^2^(ℝ), and observe that *G* is finite for each *k*, continuous in *k*, and converges to zero as *k* ↦ +∞. For each **h** ∈ *L*^2^(ℝ), we deduce that sup_*k*≥*γ0*_ *G*(*k*) < ∞, and from the uniform boundedness principal we conclude that the operators 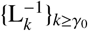 are uniformly norm bounded from *L*^2^(ℝ) to *H*^2^(ℝ). These bounds are independent of *ε* ∈ (0, *ε*_0_) as the operators are independent of *ε*. Since

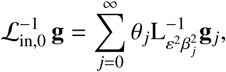

and since the Laplace-Beltrami eigenmodes are orthonormal in the *L*^2^(**S**_∞_) inner product, we deduce the existence of *M* > 0 such that (3.37) holds, and hence (3.36) follows. Since 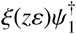 and 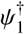 are *ε*-exponentially close, the coercivity in 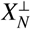 follows with an *ε* −exponentially small modification to *M*^−1^ > 0.

*Step 3: Coercivity of* ℒ_in,0_ *on* 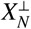 *in L*^2^(Ω). The decomposition (3.26) provides the lower bound

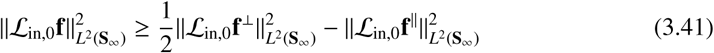

By the coercivity of ℒ_in,0_ in *Step 2*, the first positive term has a *H*^2^(**S**_∞_) lower bound while the negative term can be bounded by *H*^2^(**S**_∞_)-norm of **f**^‖^. More precisely,

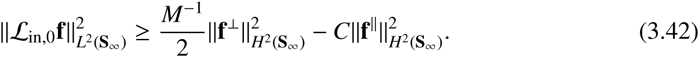

On the other hand, a second application of (3.26) yields

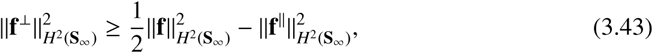

which combined with the previous inequality implies

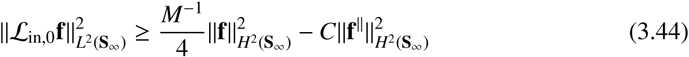

for a different constant *C*. The coercivity Lemma follows from (3.35) by replacing the *H*^2^(**S**_∞_) bound of **f**^⊥^ with the *L*^2^(**S**_∞_) bound of **f**. □

To derive a lower bound on the inner bilinear form we account for the lower order terms. Since the support of **v**_−_ lies in Γ_2ℓ_ we may apply the decomposition (3.15)

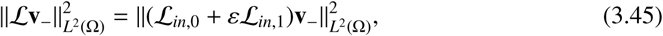

expand the quadratic form, and apply Young’s inequality to the sign-undeterminate term,

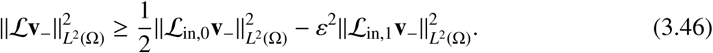

Since the support of **v**_−_ lies inside Γ_2ℓ_ its *L*^2^(Ω) and *L*^2^(**S**_∞_) norms satisfy the *ε*-equivalency of (3.24). Applying the *L*^2^-coercivity of Lemma 3.7, we arrive at the lower bound

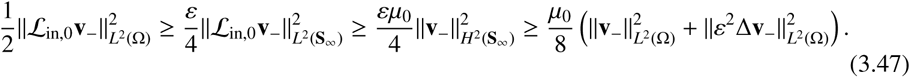

In addition, in light of the definition (3.15) of ℒ_in,1_,

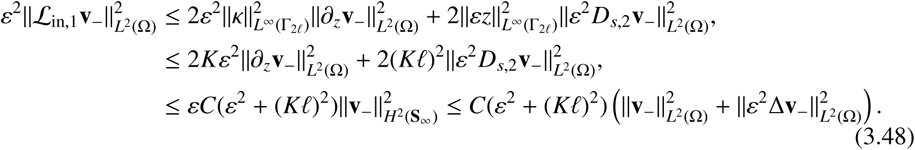

Here we used the Gargliardo-Nirenberg embedding inequality and the *ε*-equivalence of the *H*^2^(Ω) and *H*^2^(**S**_∞_) norms. Combining (3.46)-(3.48) and taking *ε* and *K*ℓ sufficiently small yields the inner coercivity:

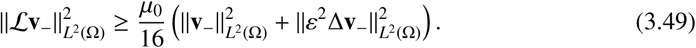

#### Outer bilinear term

Recalling the leading order, constant coefficient outer form ℒ_out,0_ of ℒ, given in (3.16), we define the outer operator 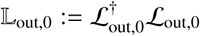.

##### Lemma 3.8.

*For any* **f** ∈ *H*^2^(Ω), *there exist µ*_0_ > 0 *such that*

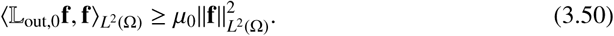

*Proof.* By assumption **a** = 0 is a normally hyperbolic equilibrium of F, in particular from Lemma 2.1 we know that *σ*(*D*^−2^ ∇_**u**_ *F*(0)) has no eigenvalues in (− ∞, 0]. Form the form of 𝕃_out,0_ we have ker(𝕃_out,0_) = ker(ℒ_out,0_), and since Ω is a rectangular box subject to periodic boundary conditions, the kernel of ℒ_out,0_ is comprised of functions of the form *e*^*ik*·*x*^**U**, where **U** ∈ ℝ^*d*^ is a constant vector. This function lies in the kernel only if

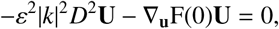

or equivalently

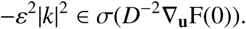

However this is precisely the condition precluded by the assumption that **a** = 0 is normally hyperbolic, thus we deduce that 𝕃_out,0_ has no kernel and is invertible. However we can make a stronger statement. The spectrum of 𝕃_out,0_ is discrete, but lies on the curves of essential spectrum defined as the finite family of dispersion relations *λ* = *λ*(*k*) for which the *d* × *d* matrix

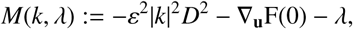

has a kernel, see chapter 3 of [19]. The assumption of normal hyperbolicity implies that none of the dispersion relation curves pass through the origin. Indeed, rescaling *k* ∈ ℝ^*d*^, the dispersion relation curves can be made independent of *ε*, and tend to −∞ as | *k* | ↦ ∞. This implies that the curves lie a finite distance *µ*_0_ > 0 to the origin, which is wholly independent of *ε*. Since 𝕃_out,0_ is self-adjoint and non-negative, this spectral bound provides the coercivity estimate (3.50). □

We expand the outer bilinear term as

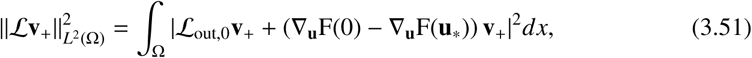

which from Young’s inequality enjoys the lower bound

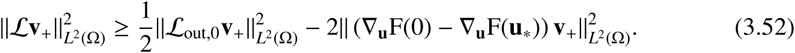

Since the support of **v**_+_ lies in Ω \ Γ_ℓ_, we have the bound

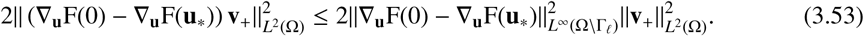

The homoclinic connection **u**_*_ converges to 0 with an exponential rate as *z* goes to infinity and the function F is smooth, so the *L*^∞^-norm of the difference on Ω \ Γ_2*l*_ is *ε*-exponential small. This establishes the bound

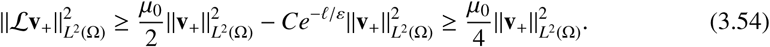

Here the constant *C* depends only on F and the decay rates of **u**_*_ in *z*.

#### Mixed bilinear terms

The support of **v**_+_**v**_−_ is contained in the overlap region Γ_2ℓ_ \ Γ_ℓ_. On this set the difference ∇_**u**_F(**u**_*_) − ∇_**u**_F(0) is *ε*-exponentially small, and we use the outer expansion of ℒ to obtain

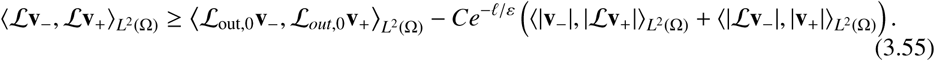

Applying Hölder’s inequality, the negative term on the right hand side can be bounded by

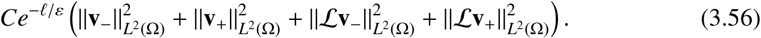

Since **v**_−_ = *ξ*(*εz*)*v*(*x*), its support is contained with Γ_2ℓ_ and we may use the inner expression for the Laplacian to obtain a lower bound on the first term on the right-hand side of (3.55). Moreover *ξ* is slowly varying in *x* and independent of *s*. With these observations we obtain the expansion

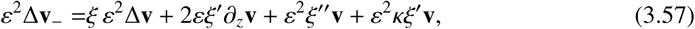

with a similar expansion for **v**_+_ with *ξ* replaced with 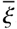. Since *ξ* and its derivatives are uniformly bounded, independent of *ε*, we obtain

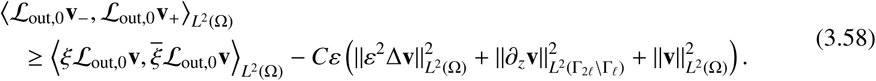

The first term on the right-hand side of (3.58) is positive since the product 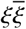 is non-negative. Using the *ε*-equivalence of *L*^2^(Ω) and *L*^2^(**S**_∞_) norms from (3.22), and standard embedding inequalities, we obtain

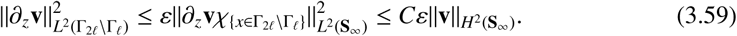

Finally, from the *H*^2^-norm *ε*-equivalence given in (3.24) we deduce

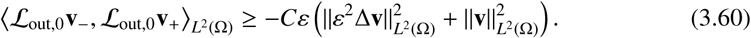

Combining the lower bounds on the inner, (3.49), outer (3.54), and mixed (3.60) bilinear forms, with the decomposition (3.20), we obtain the existence of 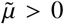, independent of *ε* for which

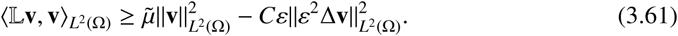

From the form of 𝕃, elliptic regularity theory affords the existence of *γ* > 0, independent of *ε* > 0, such that

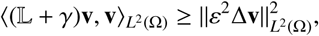

for all **v** ∈ *H*^2^(Ω). Then for any *t* ∈ (0, 1) we may interpolate

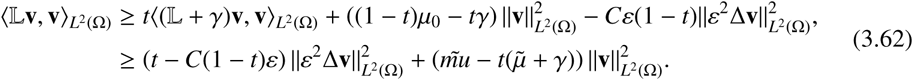

The choice 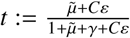 yields the estimate (3.18) with

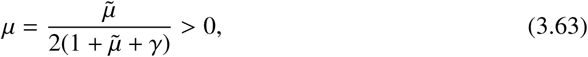

independent of *ε*. □

### 3.3 Normal coercivity of singular homoclinic freeway manifolds

The results of section 2.2 provide constructive conditions for the existence of homoclinic freeway connections in the freeway system (1.5) with F as in (2.10). Section 3.1 constructs the corresponding freeway manifold, which from Proposition 3.4 is comprised of low energy functions. Theorem 3.5 of section 3.2 equates the normal coercivity of the associated freeway manifold to a spectral condition on the linearization (3.10) of the one-dimensional freeway system at the underlying homoclinic freeway connection, **u**_*_. For the singularly perturbed systems of section 2.2, the spectral problem has been been analyzed in detail [11]. In particular the stability hypothesis of Theorem 3.5 can be related to simple geometric conditions arising in the construction of the slow-fast homoclinic freeway connections.

Assume the framework of section 2.2 and that the function ρ, defined in (2.14), has a simple root *s*_*_ > 0. Let **u**_*_ be the associated slow-fast homoclinic freeway connection. Then, under the assumption that

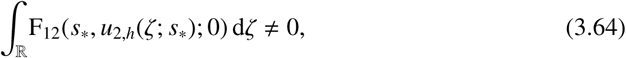

[11, Corollary 5.10 and eq. (5.16)] imply that the kernel of L, and hence that of *D*^−2^L, is simple and spanned by the translational eigenmode ∂_*z*_**u**_*_. To apply Theorem 3.5 it remains to verify that *σ*_*p*_(*D*^−2^L) has no strictly positive elements. To this end it is convenient to consider the point spectrum of the operator pencil *D*^−2^ (L − *λ*) for *λ* ∈ C. For any *k* ∈ *σ*_*p*_(*D*^−2^ (L − *λ*)) there exists a solution *Ψ* ∈ *L*^2^(ℝ) to the eigenvalue problem

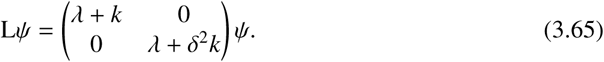

This eigenvalue problem has precisely the same structure as that in [11, eq (3.2)], modulo the replacement of ‘*λ*’ by ‘*λ* + *k*’ in the first component and ‘*λ*’ by the asymptotically close value ‘*λ* + *δ*^2^*k*’ in the second component. All the assumptions of [11] hold for this extended problem, as do each of the steps of the subsequent analysis. Indeed, the set-up of this situation is has similarities to the stability analysis of homoclinic stripes in singularly perturbed reaction-diffusion systems conducted in [26] with the exception that the case *k* > 0 was not considered therein. It follows from the prior analysis that there exists an extended analytic Evans function (*λ, k, δ*) whose roots coincide with the point spectrum of the operator pencil *D*^−2^(*L – λ*), including multiplicity. Moreover, there exists an analytic fast transmission function *t* _*f*,+_ and a meromorphic slow transmission function *t*_*s*,+_ such that the extended Evans function admits the slow-fast decomposition

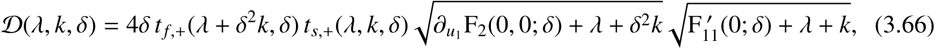

see [11, eq. (4.4)]. This Evans function decomposition, which follows from the strong structural similarity between the eigenvalue problem (3.65) and the stability problem studied in [11], allows us to prove the following Theorem.

#### Theorem 3.9.

*Suppose that the vector field* F(**u**; *δ*) *is as given in* (2.10), *the assumptions of section 2.2 hold, and s*_*_ *is a simple root of* ρ *given in (2.14). Let* **u**_*_ *be the associated freeway homoclinic connection to* **a** = 0. *Suppose that, in addition*,

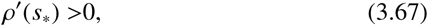

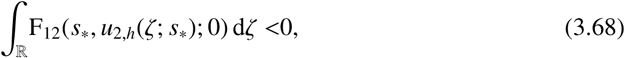

*where u*_2,*h*_(*ζ*; *u*_1_) *is as defined in Assumption 2.3. Then, the set σ*_*p*_ (*D*^−2^L)∩ℝ_+_ *consists of precisely one simple eigenvalue at the origin.*

*Proof.* The assumption (3.68), together with the fact that *s*_*_ is a simple root of ρ, guarantees the simplicity of the eigenvalue at zero. Hence, it is sufficient to show that the Evans function 𝒟(0, *k, δ*) (3.66) has no zeroes for *k* > 0 and *δ* sufficiently small. By [11, Lemma 4.3], the roots of the fast transmission function *t* _*f*,+_(*λ, δ*) are to leading order in *δ* given by the eigenvalues of the fast Sturm-Liouville operator 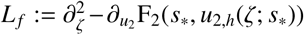. Since *L*_*f*_ is the linearization of (2.13) at the planar homoclinic *u*_2,*h*_, it has a kernel associated to the translational invariance of the planar system. This kernel is isolated and simple by the Sturm separation theorem. Hence, by the inverse function theorem, *t* _*f*,+_(*δ*^2^*k, δ*) ≠ 0 for sufficiently small *δ*.

By [11, Theorem 4.4], we can express the slow transmission function *t*_*s*,+_(*λ, k, δ*) to leading order in *δ* as

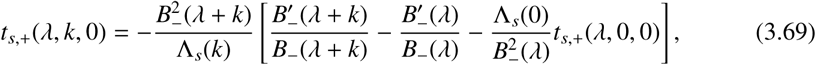

where 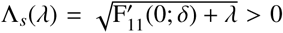 (cf. [11, eq. (3.8)]) and *B*_−_(*λ*), *B*′_(*λ*) are as defined in [11, Theorem 4.4]. By [11, Lemma 5.9], for *λ* = 0, we can write

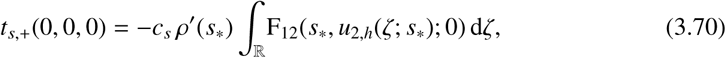

with *c*_*s*_ > 0, using [11, eq. (2.9)]. From [11, Lemma 5.6] we know that *B*_−_(*λ*) ≠ 0 for all *λ* ≥ 0 if and only if *y*_*_ > 0, where

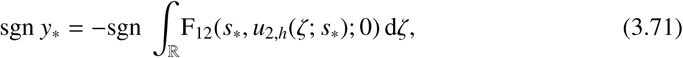

see [11, Lemma 2.2]. We employ a Prüfer transformation [11, eq. (5.5)] to write

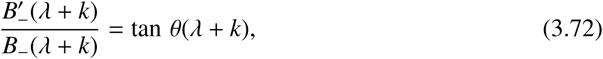

where θ : ℝ ↦ ℝ. From the statement of [11, Lemma 5.4] we deduce the strict monotonicity of θ, and conclude

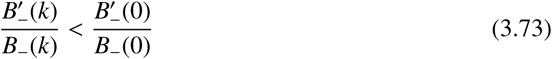

for all *k* > 0. Combining (3.70) with the assumptions (3.67) and (3.68) implies that

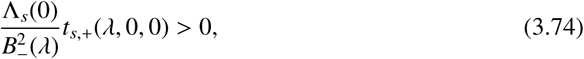

which can be taken together with (3.73) to conclude that factor within the square brackets in (3.69) is negative, while the prefactor *B*_−_(*λ* + *k*) is finite and never zero. We deduce that *t*_*s*,+_(0, *k*, 0) > 0 for all *k* > 0. The non-vanishing of the Evans function (3.66) for *λ* = 0 now follows from [11, Corollary 4.2]. □

#### Corollary 3.10.

*Suppose that the assumptions of Theorems 3.5 and 3.9 are met. Then, there exists δ*_0_, 𝒢_0_>*0 for which each δ* ∈ (0, *δ*_0_) *and each K*, ℓ > 0 *satisfying K* ℓ < 𝒢_0_ *yield an ε*_0_ > 0 *and a µ* > 0 *such that the freeway homoclinic connection* **u**_*_ *of* (1.5) *corresponding to the system presented in* (2.10) *generates a normally coercive manifold* ℳ_*K*,ℓ_ (**u**_*_), *satisfying (3.18) with coercivity constant µ for all* 𝕃 = L_Γ_ *with* Γ ∈ 𝒢_*K*,ℓ_.

The PCB system presented in section 2.3 prescribes a take-off curve and an unstable slow manifold, as depicted in Figure 3. When the take-off curve crosses the unstable manifold from above, as it does at *u*_1_ = *s*_*_, then ρ′(*s*_*_) > 0 and Corollary 3.10 holds. In particular the freeway manifold generated by **u**_*_ is normally coercive in the sense of Theorem 3.5.

## 4 Freeway to Toll-Road Bifurcations

Minimizers of the reduced free energy (1.4) solve the toll-road system (1.6). In this section we consider bifurcations within the freeway system (1.5) that induce changes in solution type within the larger toll-road system. We insert a parameter, *µ*, within the vector field F = F(·; *µ*). When written as pair of second order systems, the toll-road system (2.1) has the equivalent formulation

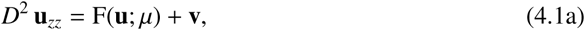

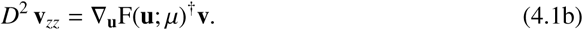

The freeway solutions satisfy (4.1) with **v** = 0.

In this section, we assume that for *µ* ≥ 0 the toll-road system (4.1) admits a one-parameter pair of freeway connections (**u**_±_(*µ*), 0)^*t*^ between the fixed equilibria **a**_*i*_ and **a** _*j*_. Moreover, we assume that the two branches merge at *µ* = 0 through a saddle-node bifurcation, with **u**_+_(0) = **u**_−_(0) := **u**_0_. We shift the origin (**u, v**) ↦ (**u**_0_ + **u, v**) and expand (4.1) around the connection (**u**_0_, 0)^*t*^ at the saddle-node bifurcation *µ* = 0. This results in the formulation

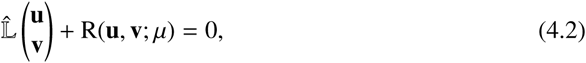

where we have introduced

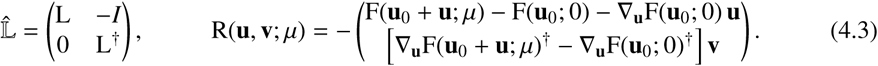

As before L, defined in (3.10), is the linearization of (1.5) at **u**_0_ with *µ* = 0. The nonlinear remainder term ℝ(**u, v**; *µ*) (4.3) can be expanded for small (**u, v**)^*t*^ and small *µ*, yielding

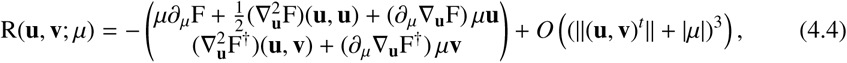

where we assume for simplicity that F(**u**; *µ*) depends linearly on µ, i.e. 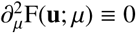.

We assume that the saddle-node bifurcation at *µ* = 0 is non-degenerate. Due to translational invariance *Ψ*_1_ := ∂_*z*_**u**_0_ ∈ ker(L) and the saddle-node bifurcation yields another central direction. Specifically

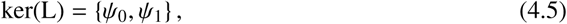

with

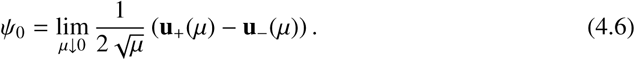

From the structure of 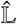 and the Fredholm alternative we deduce that

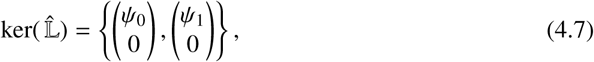

where *Ψ*_0_ and *Ψ*_1_ are even and odd, respectively, about *z* = 0. We introduce ker 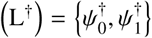, with 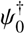 and 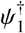 also even resp. odd about *z* = 0. The spectral projections onto *Ψ* _*j*_ and 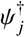 are given by

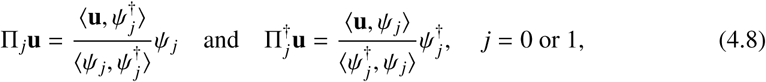

with complementary projections 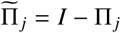 and 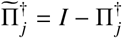.

### 4.1 Normal form expansion

We perform a normal form expansion in (4.2). We write the perturbative term (**u, v**)^*t*^ in the form

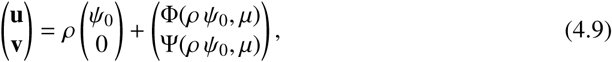

where the nonlinear functions Φ, Ψ are expanded as

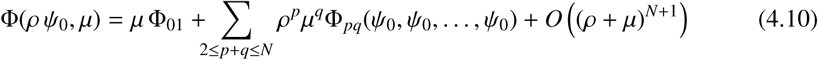

for small *ρ* and *μ*; here, Φ_*pq*_ is a *q*-linear map. Ψ is expanded analogously.

#### *Remark* 4.1.

While the translational invariance of (1.6) introduces a central direction through the *z*-derivative of **u**_0_, the same translational invariance precludes *Ψ*_1_ = ∂_*z*_**u**_0_ to play a direct role in the normal form expansion (4.9). This is a direct consequence of [15, Theorem 3.19], see also [29, Theorem 3.3]. Hence, (4.9) does not contain a linear term of the form 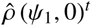, nor do the nonlinear functions Φ and Ψ explicitly depend on 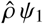.

Substitution of the normal form expansion (4.9) in (4.2) yields at *O*(*μ*)

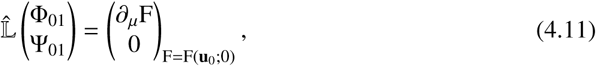

which by the definition of 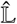 (4.3) is equivalent to

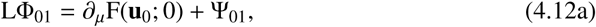

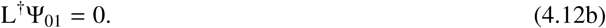

We see that Ψ_01_ ∈ ker(L^†^); hence, the solvability condition of (4.12a) yields

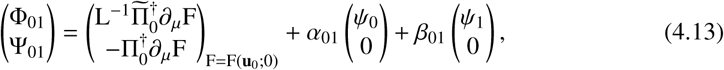

with *α*_01_ and β_01_ yet to be determined. Next, we consider the equation at *O*(*μ*^2^)

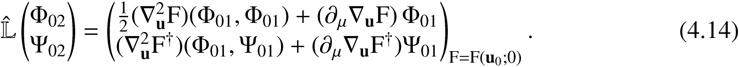

The solvability condition for the equation for Ψ_02_ stipulates that

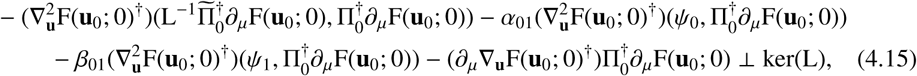

from which follows that *β*_01_ = 0 and

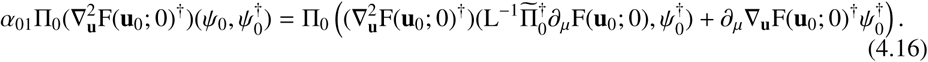

The equation at *O(ρ*^2^)

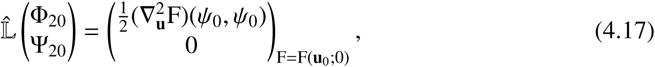

being of the same qualitative form as (4.11), can be solved to obtain

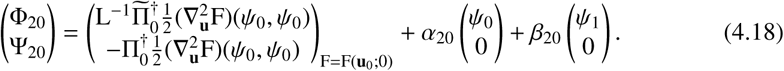

However, the equation at *O*(*ρμ*)

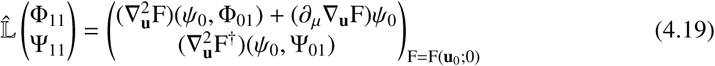

yields as solvability condition for Ψ_11_

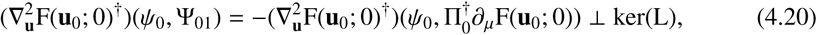

which is in general *not* satisfied. At the next order, we encounter a similar situation at *O*(*ρ*^3^), where the equation

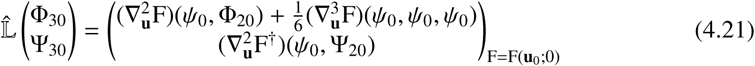

yields as solvability condition for Ψ_30_

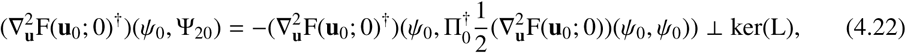

which is also in general not satisfied. Furthermore, the equations at *O*(*ρμ*^2^) and *O*(*ρ*^2^*μ*) explicitly depend on Ψ_11_, the term that yielded the problematic solvability condition (4.20).

To resolve these issues, we assume a *resonance* for the problematic equations at *O*(*ρμ*) and *O*(*ρ*^3^) [15]. We take *p, q* ∈ ℤ_≥0_, *p* + *q* ≥ 1, such that *ρμ* = *ρ*^*p*^*μ*^*q*^; likewise, we assume that there exist *r, s* ∈ ℤ_≥0_, *r* + *s* ≥ 1, such that *ρ*^3^ = *ρ*^*r*^*μ*^*s*^. From these assumptions, it follows that

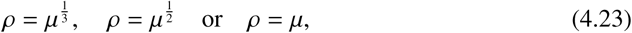

where we ruled out *ρ* = *μ*^*k*^ with *k* > 1, by standard arguments. The choice 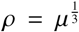 yields the same insolvable equation at *O*(*μ*) = *O*(*ρ*^3^) while the choice 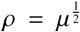 yields a transverse bifurcation with persistence of the freeway solutions for *μ* > 0. Hence, the only relevant scaling choice to be investigated is *ρ* = *μ*.

To simplify notation, we rewrite the normal form expansion (4.9), (4.10) and set

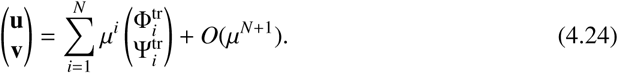

Substitution of the normal form expansion (4.24) in (4.2) yields at *O*(*μ*)

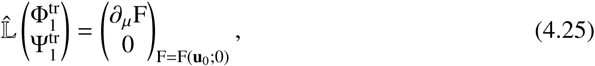

which is equivalent to (4.11); hence, we obtain

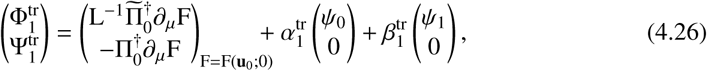

with 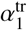 and 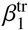 to be determined at the next order. At *O*(*μ*^2^), we find

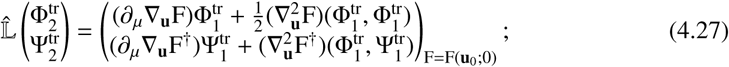

the solvability condition for 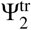 yields 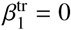 and

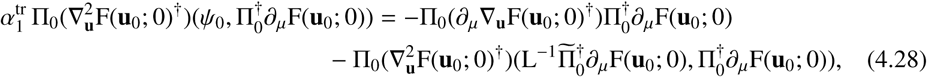

which fully determines 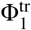 (4.26). Furthermore, we obtain

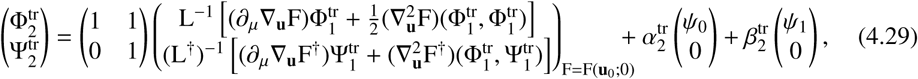

with 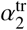 and 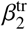 to be determined at the next order. This expansion allows us to formulate the following Theorem:

#### Theorem 4.2.

*Let* 0 < *δ* ≪ 1 *be sufficiently small. Assume that there exists μ*_0_ > 0 *such that the freeway system* (1.5) *admits a pair of orbit families* **u**_±_(μ) *connecting the same equilibria* **a**_*i*_ *and* **a** _*j*_ *for all* 0 < *μ* < *μ*_0_; *assume that this pair of orbit families coincides and terminates at* **u**_+_(0) = **u**_−_(0) = **u**_0_ *through a saddle-node bifurcation; assume that this saddle-node bifurcation is nondegenerate. Denote the linearization of* (1.5) *at* **u**_0_ *by* L (3.10). *Then, there exists an open neighbourhood U of μ* = 0 *such that for all μ* ∈ *U, there exists a minimizer* **u**_tr_(*μ*) *of the reduced free energy* ℱ_1_ (1.4), *with energy value*

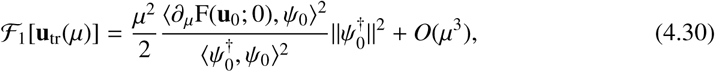

*with Ψ*_0_ *as in* (4.6), *and where* 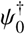 *is the unique element of ker*(L^†^) *that is even as a function of z.*

*Proof.* The local existence of **u**_tr_(*μ*) for small *μ* is an immediate consequence of the normal form expansion in section 4.1. The reduced free energy (1.4) can be written in terms of the norm induced by the *L*^2^(ℝ)-inner product as 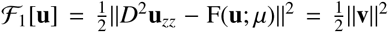 by (4.1a).The leading order expansion of **v** given in (4.26) yields the energy value to leading order in *μ.* □

#### *Remark* 4.3.

The existence of homoclinic orbits in (1.5) as presented in [11] is a consequence of the transversal intersection of manifolds, which is directly equivalent to the invertibility of L (3.10) (up to translation). This implies invertibility of 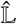 (4.3), and ensures the unique local embedding of solutions of (1.5) in the phase space of (4.1). The toll-road branch **u**_tr_ that intersects the freeway homoclinic families **u**_±_ at *μ* = 0, exists precisely because the invertibility of L fails at *μ* = 0, introducing a nontrivial (even) kernel element *Ψ*_0_, which is the basis for the normal form expansion in section 4.1.

### 4.2 Toll-road connections in the PCB model

The bifurcation analysis of section 4 allows the construction of low-energy toll-road connections. This is relevant to situations in which mass constraints prevent the formation of freeway connections. For the PCB model of section 2.3, the results of Theorem 4.2 can be applied by extending the take-off curve to depend upon the bifurcation parameter *μ*, that is *T*_o_(*s*) = *T*_o_(*s*; *μ*). In particular we make the following assumptions.

#### Assumption 4.4.

*Let T*_o_(*s*) = *T*_o_(*s*; *μ*) *depend on a parameter μ, and let* ρ_PCB_ = ρ_PCB_(*s*; *μ*) *accordingly be as in* (2.19). *There exists s*_sn_ ∈ (0, *u*_1,max_) *for which* 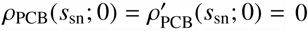 *and*

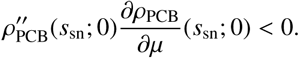

These assumptions guarantee the local existence of a pair of families of homoclinic orbits in the freeway system (1.5) that terminates in a nondegenerate saddle-node bifurcation when *μ* = 0. For the PCB model (2.16), we find

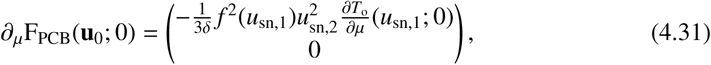

where **u**_sn_ = (*u*_sn,1_, *u*_sn,2_)^*t*^ is the (degenerate) homoclinic orbit at the saddle-node bifurcation. Using Theorem 2.5, we can obtain an explicit expression for *Ψ*_0_ as defined in (4.6), as follows. From Assumption 4.4, it follows that the pair of solutions *s*_±_(*μ*) to ρ_PCB_(*s*; *μ*) = 0 can be expanded as 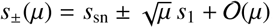, with

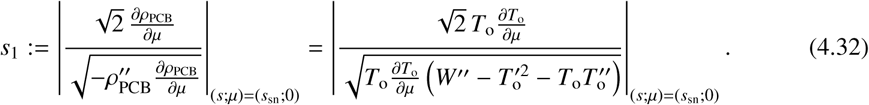

Moreover, writing 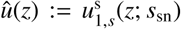 (for the definition of 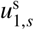, see Theorem 2.5), we see that there exists a shift *z*_1_ < 0 such that 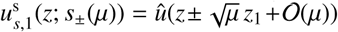; a direct calculation shows that 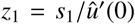. Hence, the saddle-node eigenvector *Ψ*_0_ has, by Theorem 2.5, the following leading order structure:

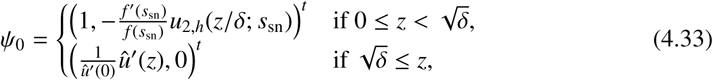

where *Ψ*_0_ has been scaled by *s*_1_ compared to its original definition (4.6). We now use (4.33) to calculate

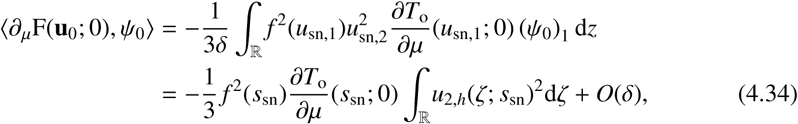

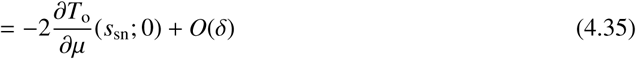

by (2.17). Furthermore, we know that 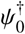 is the unique even element of ker L^†^, which therefore solves the system

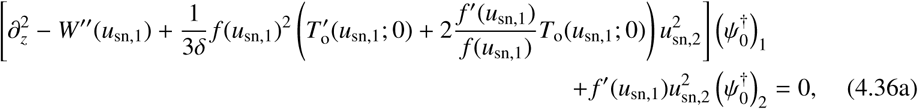

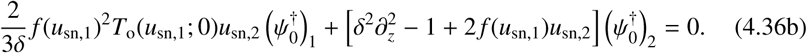

This system can be significantly simplified using the scale separated structure of the underlying homoclinic **u**_sn_ as given in Theorem 2.5. In particular, outside the symmetric interval 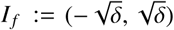, system (4.36) reduces to

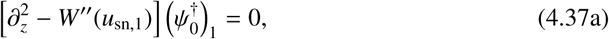

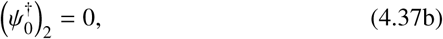

up to *δ*-exponentially small terms. We note that 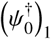 must be a multiple of 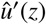, and 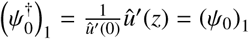 without loss of generality. Inside *I*_*f*_, we rescale *ζ* = *z*/*δ* and find

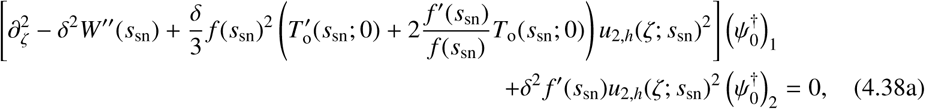

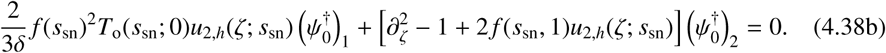

From (4.38b), we infer that 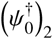 scales with 1/*δ*. For the first component 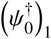, this yields 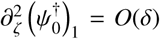 from which we conclude 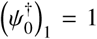 by continuity. Rescaling the second component 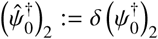, it obeys

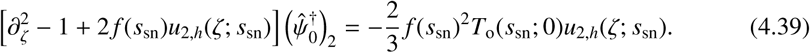

Using (2.17), we can reduce (4.39) to

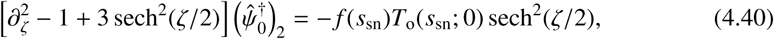

which can be solved explicitly, yielding

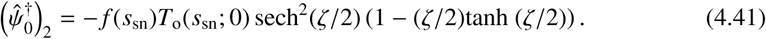

To summarize, we have found to leading order in *δ*

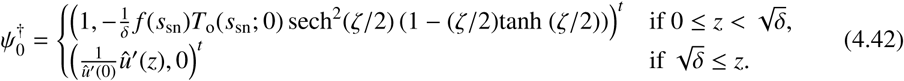

This allows us to calculate

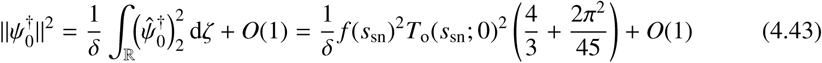

and

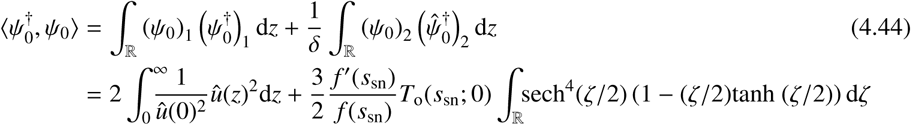

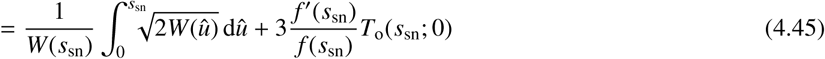

to leading order in *δ*. Using the results obtained so far, we calculate the value of the reduced free energy of the toll-road branch in the PCB model:

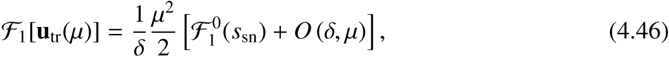

with

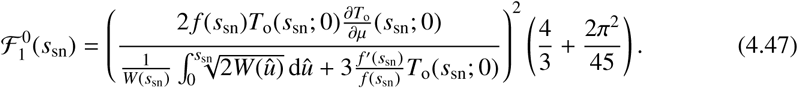

For a PCB model with a prescribed take-off curve, embedding the take-off curve in a larger familty *T*_o_(*s, μ*) which has a saddle-node bifurcation at *μ* = 0 and reverts to the original take-off curve at *μ* = *μ*_*_, provides for the existence of a toll-road connection with cholesterol mass scaled by *f* (*s*_sn_) with energy given by (4.46) with *μ* = *μ*_*_. This relates the distance of the take-off curve to the unstable slow manifold to the existence and energy of an associated toll-road connection.

